# Long-term structural plasticity of hippocampal dendritic spines following contextual fear memory reactivation

**DOI:** 10.1101/2023.05.30.542970

**Authors:** Candela Medina, Santiago Ojea Ramos, Lucas Pozzo Miller, Arturo Romano, Verónica de la Fuente

**Affiliations:** Laboratorio de Neurofarmacología de los Procesos de Memoria, Cátedra de Farmacología, Facultad de Farmacia y Bioquímica, Universidad de Buenos Aires (UBA), Buenos Aires, Argentina; Instituto de Fisiología, Biología Molecular y Neurociencias (IFIBYNE-UBA-CONICET), Buenos Aires, Argentina; Department of Neurobiology, University of Alabama at Birmingham, Birmingham, Alabama, USA; Departamento de Fisiología, Biología Molecular y Neurociencias, Facultad de Ciencias Exactas y Naturales, Universidad de Buenos Aires, Buenos Aires, Argentina

**Keywords:** dendritic spines, spine morphology, dorsal hippocampus, fear conditioning, memory, reconsolidation, NF-κB

## Abstract

Dendritic spines are plastic structures exhibiting a high degree of morphological variability. Certain morphometric parameters, such as volume, positively correlate with the strength of the synapse in which they participate. Memories, too, are subject to change over time and with experiences. In particular, the presence of a reminder of a learning event can trigger the labilization of the memory trace, followed by a re-stabilization process termed reconsolidation. The underlying mechanisms behind the labilization/reconsolidation processes are of great interest, as they are thought of as possible targets for the treatment of post-traumatic stress disorders. Dendritic spines have long been considered the physical sites for memory formation and storage. Our work aimed at studying the long-term spine morphological plasticity associated with labilization/reconsolidation in the dorsal hippocampus, a brain region relevant for the formation of contextual memories. Our results suggest that labilization/reconsolidation does not affect spine density, but rather induces changes in spine morphology. Furthermore, we show that some of these changes are prevented by the inhibition of the transcription factor NF-κB inhibition. Finally, we found that NF-κB negative modulation also affects spine morphology in animals that were not exposed to recall but have undergone the training session, suggesting that there may be a late surge of NF-κB activity resulting from the consolidation itself.

## Introduction

Dendritic spines are small protrusions that extend from the dendrites and are the primary sites for receiving synaptic inputs from other neurons. Despite the high degree of morphological variability they exhibit, some morphological parameters are decisive in inferring the strength of the synapse in which they participate ^1–4^. In particular, the volume of the spine head is directly proportional to the size of its postsynaptic density (PSD) and the number of glutamate receptors, which determines the efficacy of synaptic transmission^3,5–7^. In addition, the area of the PSD is directly proportional to the area of the active zone on the presynaptic side, as well as to the number of docked vesicles and the total number of vesicles per bouton^8^, a reliable indicator of the amount of neurotransmitter released per action potential^9^. Thus, large head spines are the sites of strong synapses in terms of both postsynaptic sensitivity and neurotransmitter release. Dendritic spine necks, which connect the spine head to the parental dendrite, are also postulated to play an important role in synaptic efficacy, modulating electrical coupling, calcium exchange and diffusion of signaling molecules between the spine head and the parental dendrite^10–13^.

Since their early discovery by Ramon y Cajal in 1888^14^, dendritic spines have been postulated as the anatomical loci for memory formation and storage. First, they have been demonstrated to undergo changes in size, density and shape relatively quickly^15,16^, but are also able to maintain their structure for years ^17,18^. These two characteristics are compatible with both the speed of a learning event and with the long lasting feature of many memories. Second, *ex vivo* long-term potentiation (LTP), an enduring enhancement of synaptic transmission resulting from particular types of electrophysiological stimulations which is proposed to be a cellular correlate of learning and memory^19^, induces structural changes in spines^20–25^. Indeed, it has been shown that LTP at individual spines using two-photon photolysis of caged glutamate induces their enlargement. Interestingly, the temporal stability of the changes depended on the size and shape of the spine prior to stimulation, being transient in large mushroom-like spines but persistent in small spines^16^. Third, *in vivo* studies in which individual spines were imaged over time demonstrated that new experiences increase the turnover of spines, favoring both spine pruning and spine formation, and that many newly formed spines, as well as already present spines, remain stable over time^26^.

For many years, efforts have been devoted to understanding the structural changes in neural circuits involved in learning and memory-related processes, even in models of memory that rely on deep brain areas where it is methodologically difficult to study individual spines over time *in vivo*^25,18,4^. A widely used paradigm to study mechanisms underlying memory formation fear conditioning, which is also of particular interest because its study allows for a better understanding of the physiological foundations of fear-related dysfunctions in humans, such as phobias and post-traumatic stress disorders^27,28^. It has been well established that the presentation of a reminder of a learning event can trigger the reactivation of that memory trace^29,30^. Reactivation gives rise to a period in which the trace becomes labile, being susceptible to disruption. A process known as reconsolidation is needed to re-stabilize the trace^31–33^. There is general consensus that the stabilization of fear memories over time, *i.e*. their consolidation, involves structural plasticity^4,34^. However, there are very few studies investigating the structural plasticity associated with experience-dependent modifications of already established fear memories, particularly those that result in the stabilization/reconsolidation of a fear memory^35–37^. Indeed, to our knowledge, there are no reports on changes in morphometric parameters of dendritic spines associated with labilization/reconsolidation processes. The aim of this study was to investigate whether the reactivation of a memory trace leads to long lasting remodeling of dendritic spines. To achieve this, we used the context-fear conditioning paradigm, and focused on the dorsal CA1 region of the hippocampus, a brain area known to be involved in the memory processes elicited by contextual fear memory reactivation^38^. In addition, for a mechanistic and functional study, we chose to modulate memory reconsolidation using pharmacological inhibition of the transcription factor NF-κB (nuclear factor κB)^39–42^.

## Results

### 1. Reconsolidation does not affect total spine density in either apical or basal dendrites of CA1 pyramidal neuron of the dorsal hippocampus

The process of reconsolidation allows the eventual strengthening of a memory trace^43^, and/or the incorporation of new information related to a pre-existing one^44,45^. However, in general, in many experimental conditions the only way to demonstrate its existence is by blocking it. Here, our aim was to study long-term morphological changes elicited by the reactivation of the memory trace, or in other words, its labilization/ reconsolidation. Considering the possibility that these changes might be transient, we decided to include a pharmacological strategy known to disrupt the reconsolidation of hippocampal dependent contextual fear memories: the blockade of the transcription factor NF-κB^39–42^. We hypothesized that this tool would help us to assess long term effects, should they exist, at a morphological level as well. The strategy consisted of injecting in the dorsal hippocampus a double-stranded DNA oligonucleotide containing its consensus sequence (κB Decoy, Dec from now on)^46,47,40,41^. This oligonucleotide enters the cells and induces NF-κB inhibition 15 min after intrahippocampal injection^48^. A mutated Decoy oligonucleotide (mDecoy, mDec from now on) was used as a control for a possible nonspecific effect of DNA administration in the hippocampus. mDec is a stringent control because the entire overall composition of bases is conserved, except for one base in the consensus sequence that is mutated, impeding NF-κB binding^47,48^. We first checked that NF-κB inhibition in the dorsal CA1 region indeed blocks reconsolidation. For this, we trained cannulated mice, and 24 h later we re-exposed them for 5 min to the training context, a reminder that elicits labilization and reconsolidation of contextual fear memories^41,49–51^. Animals were divided into two groups, one which was injected intrahippocampally with the NF-κB inhibitor (RE-Dec). The other received the control solution (RE-mDec). Memory was assessed 48 h after training (Fig. 1A,B). Only the RE-Dec group exhibited a long-term memory deficit on day 3, demonstrating that NF-κB inhibition disrupted memory reconsolidation (Fig. 1C; RE-Dec vs RE-mDec: p = 0.0003).

**Figure 1.**
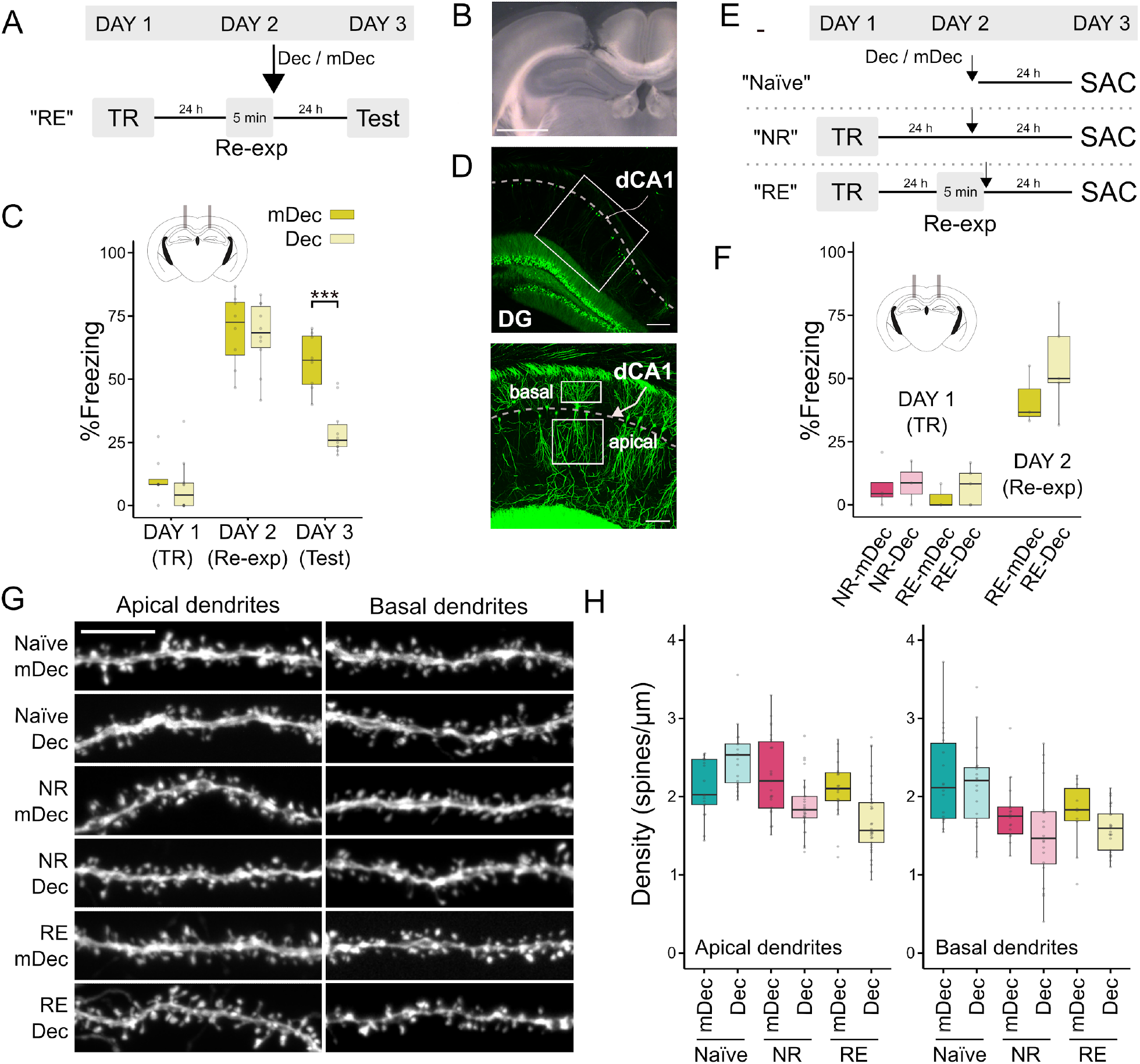
Reconsolidation does not affect total spine density in either apical or basal dendrites of CA1 pyramidal neuron of the dorsal hippocampus. **(A)** Contextual fear conditioning design: Four groups of cannulated mice were trained in a fear conditioning task. Twenty four hours later, two groups were re-exposed for 5 min to the training context, one of them was injected with decoy (RE-Dec), and the other with mDec (RE-mDec). The two remaining trained groups were injected with Dec or mDec but not re-exposed to the training context (NR-Dec and NR-mDec, respectively). Memory was assessed 48 h after training. **(B)** Percentage of freezing. Data are presented as box plots, depicting the first quartile, median, third quartile. The upper whisker extends to the largest value within 1.5 times the interquartile range (IQR), while the lower whisker extends to the smallest value within 1.5 times the IQR. ^*^P < 0.05. n=8-10. pre-TR: 2 min exposure to training context to establish basal freezing level; TR: training; Re-exp: context re-exposure. **(C)** Example of a cannula above the dorsal hippocampus. Scale bar 1 mm. **(D)** Upper: Coronal section including part of the dorsal hippocampus of a thy1-GFP mouse, DG: dentate gyrus. Scale bar 200 μm. A square indicates the approximate region for the lower image. Lower: Z stack. The regions from which images were taken to study basal and apical dendrites are shown. Scale bar 100 μm **(E, F)** Same as in A, B but for thy1-GFP mice, sacrificed 48 h after TR without testing session. ^*^P < 0.05; n=3-5. n=2-5. **(F)** Same as in B but for thy1-GFP mice. **(G)** Spine density in apical (left) and basal (right) dendrites of the dorsal CA1. **(H)** Representative images of the groups. Scale bar: 5 μm. Full statistical data can be found in Suppl. Table 2.

To study structural plasticity, we used thy1-GFP transgenic mice, which present a sparse and bright Golgi-staining-like labeling of pyramidal neurons in the dorsal CA1 (Fig 1D). We were interested in assessing the morphological state of the dendritic spines at the same temporal point where memory had been evaluated in the previous experiment, *i*..*e*. 48 h after training. Thus, we repeated the experiment, but instead of behavioral testing, we perfused animals and prepared their brains for structural studies. We added two groups of behaviorally naïve animals that were intrahippocampally injected with either Dec or mDec (Na-Dec and Na-mDec, respectively). Also, we included two groups of trained mice that were injected with Dec or mDec but were not re-exposed to the training context (NR-Dec and NR-mDec, respectively) (Fig. 1E,F). Importantly, NF-κB inhibition without memory reactivation does not affect contextual fear memory^41^. The following comparisons were performed throughout the study to unravel different phenomena of interest. Firstly, from a behavioral perspective, the comparison between RE-mDec and NR-mDec will account for phenomena associated with memory stabilization/reconsolidation. In turn, the comparison between NR-mDec and Na-mDec will evaluate changes resulting from consolidation, since the only difference between the two groups is training. Secondly, regarding NF-κB modulation, the comparison between Na-mDec and Na-Dec will account for a possible basal role of NF-κB in the morphology of the spines and dendrites of interest. In turn, the comparison of R-mDec and R-Dec will assess a possible reconsolidation-dependent role of NF-κB on morphological plasticity (eventually added to possible late effects of NF-κB on consolidation). Finally, the comparison of NR-mDec and NR-Dec will account for possible roles of NF-κB in late consolidation (since the injection of its inhibitor is 24 h after training).

We analyzed apical and basal dendrites separately (Fig. 1G,H), as it has been shown that they receive different inputs ^52–54^, and differ in excitability ^55^, suggesting its differential involvement in physiologic processes^56^. Groups did not differ in their total spine density, both in apical and basal dendrites. Importantly, NF-κB inhibition did not alter total spine density in behaviorally naïve animals.

### 2. Spine density, per type of spine

According to their shape and size, spines have been commonly classified into mushroom, stubby, and thin^57,2^. We proceeded to analyze the density of spines under this classification (Fig 2, Suppl. Fig. 1).

**Figure 2.**
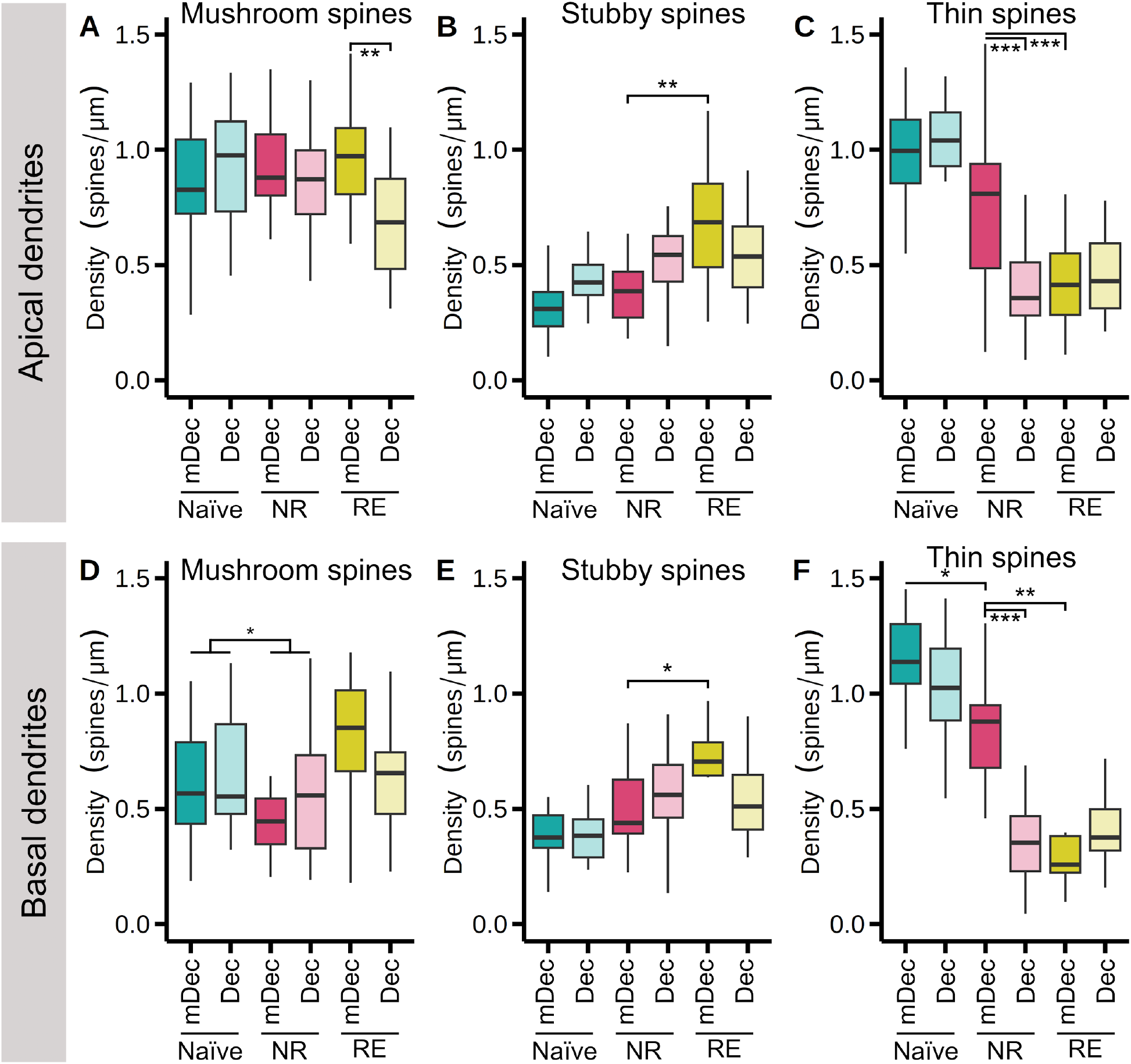
Long-term changes in spine density, analyzed per type of spine, associated with either the reconsolidation process or late-consolidation. Mushroom **(A)**, stubby **(B)** and thin **(C)** spine density of dorsal CA1 apical dendrites. **(D**,**E**,**F)** Same as in (A,B,C) basal dendrites. In all cases data are presented as box plots, depicting the first quartile, median, third quartile. The upper whisker extends to the largest value within 1.5 times the IQR, while the lower whisker extends to the smallest value within 1.5 times the IQR. n=2-5. ^*^P < 0.05, ^**^P < 0.01, ^***^P < 0.001. Full statistical data can be found in Suppl. Table 2.

#### Mushroom spine density

When analyzing the apical dendrites of dorsal CA1 (Fig 2A, Suppl. Fig. 1A) of animals injected with control solution mDec, we found that re-exposed mice did not differ in mushroom spine density when compared to non re-exposed mice (RE-mDec vs. NR-mDec: p = 0.9649). Furthermore, this variable did not present significant differences between behaviorally naïve animals and non re-exposed animals, both injected with mDec (Na-mDec vs. NR-mDec: p = 0.8907). These results suggest that neither reactivation of memory, nor training *per se* (*i.e*. trained but non re-exposed animals), have a long-term effect on total mushroom density. In turn, inhibiting NF-κB after re-exposure decreased the mushroom spine density, compared to re-exposed mice injected with mDec (RE-mDec vs. RE-Dec: p < 0.01). This decrease, together with the previous results, suggests that although re-exposure to the training context does not alter long-term mushroom spine density, there is indeed an active and dynamic change in morphology associated with memory reactivation, which is only evident when reconsolidation is altered by NF-kB inhibition. Of interest, NF-κB inhibition did not alter mushroom spine density in non re-exposed animals (NR-mDec vs. NR-Dec: p = 0.9999), nor in the behaviorally naïve mice (Na-mDec vs. Na-Dec: p = 0.8784).

In regard to mushroom spines of basal dendrites of the dorsal CA1 (Fig 2D, Suppl. Fig. 1D), we found a statistical general effect of behavior. Re-exposed mice had an increase of mushroom spine density compared to non re-exposed mice (RE vs. NR: p < 0.05).

#### Stubby spine density

In apical dendrites (Fig 2B, Suppl. Fig. 1B) of animals injected with the control solution mDec, we found that re-exposure increased stubby spine density, compared to non re-exposed animals (RE-mDec vs. NR-mDec: p < 0.01). Furthermore, we did not find an effect of NF-κB inhibition dependent on the re-exposure (RE-mDec vs. RE-Dec: p = 0.4834). Similar results were obtained when analyzing stubby spine density in basal dendrites (Fig 2E, Suppl. Fig. 2E; RE vs. NR: p < 0.05).

#### Thin spine density

In apical dendrites (Fig 2C, Suppl. Fig. 1C) of animals injected with the control solution mDec, re-exposure decreased thin spine density, compared to non re-exposed animals (RE-mDec vs. NR-mDec: p < 0.001). In addition, non re-exposed animals did not differ from behaviorally naïve animals (Na-mDec vs. NR-mDec: p = 0.9998). In turn, inhibiting NF-κB after re-exposure had no effect on thin spine density, compared to re-exposed animals injected with mDecoy (RE-mDec vs. RE-Dec: p = 0.9998). Surprisingly, NF-κB inhibition indeed had an effect on thin spine density in non re-exposed animals (NR-mDec vs. NR-Dec: p < 0.001). This latter result is interesting because it has been shown that NF-κB inhibition does not alter memory if the reminder is not presented^42,48^. Our results suggest that although the behavior we measure as an inference of memory may not be altered, the brain may indeed change with the inhibition of this nuclear factor.

When analyzing thin-type spine density of basal dendrites (Fig 2F, Suppl. Fig. 2F) of animals injected with the control solution mDec, we obtained similar results. Re-exposure decreased thin spine density compared to non re-exposed animals (RE-mDec vs. NR-mDec: p < 0.001). Here, however, we did find differences between non re-exposed animals and behaviorally naïve mice (Na-mDec vs. NR-mDec: p < 0.05), suggesting that there is a long-lasting effect of training *per se* on structural plasticity. Of interest, we again observed a decrease in thin spine density due to NF-κB inhibition in non re-exposed animals (NR-mDec vs. NR-Dec: p < 0.001).

### 3. Morphometric parameters of dendritic spines

As mentioned, spines can be classified into the categories mushroom, stubby and thin according to a set of different morphometric parameters^2^. However, within each category there is a certain degree of variability in the values of each parameter. Thus, our experimental groups could still differ in some parameters within a spine category. We next sought to investigate how similar are the mushroom, thin, and stubby spines-respectively-from each behavioral group. In order to achieve this goal, we evaluated the spines’ volume, together with their head and neck diameter.

#### i) Spine volume

##### Mushroom spine volume

The volume of mushroom spines from both apical and basal dendrites did not change either with behavior nor with NF-κB inhibition (Fig 3A,D Suppl. Fig. 2A,D).

**Figure 3.**
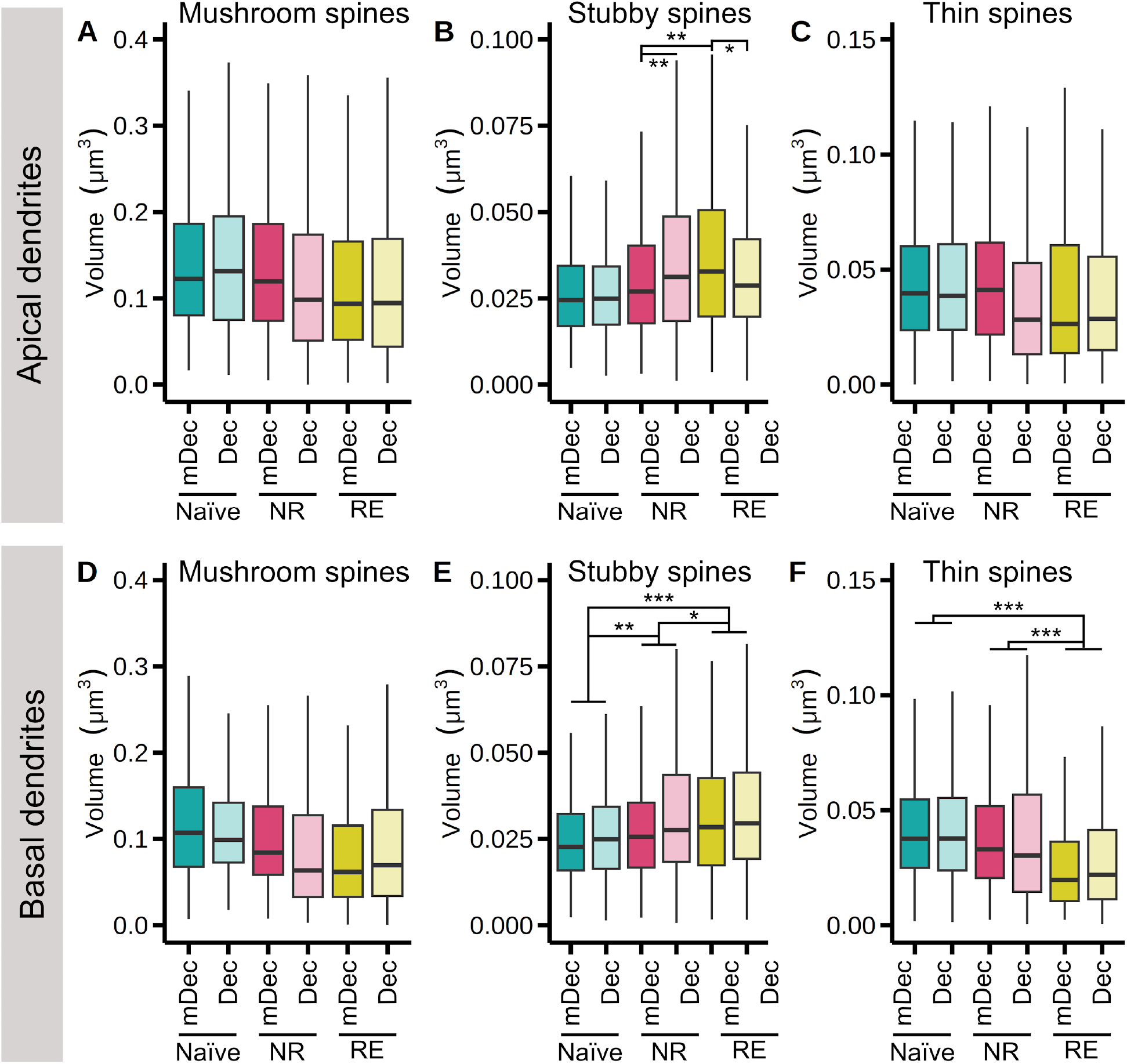
Long-term changes in volume of dendritic spines, analyzed per type of spine, associated with either the reconsolidation process or late-consolidation. Mushroom **(A)**, stubby **(B)** and thin **(C)** spine volume in the dorsal region of CA1, apical dendrites. **(D**,**E**,**F)** Same as in (A,B,C) in for basal dendrites. In all cases data are presented as box plots, depicting the first quartile, median, third quartile. The upper whisker extends to the largest value within 1.5 times the IQR, while the lower whisker extends to the smallest value within 1.5 times the IQR. n=2-5. ^*^P < 0.05, ^**^P < 0.01, ^***^P < 0.001. Full statistical data can be found in Suppl. Table 2.

##### Stubby spine volume

In apical dendrites (Fig 3B, Suppl. Fig. 2B) of animals injected with the control solution mDec, re-exposure increased the volume of stubby spines, compared to non re-exposed animals (RE-mDec vs. NR-mDec: p < 0.01). In turn, non re-exposed animals had similar levels of stubby spines’ volume compared to behaviorally naïve animals (Na-mDec vs. NR-mDec: p = 0.1733). In turn, inhibition of NF-κB after re-exposure decreased thin spine’ volume, compared to re-exposed animals injected with mDecoy (RE-mDec vs. RE-Dec: p < 0.05). Surprisingly, inhibition of NF-κB in non re-exposed animals increased the volume of stubby spines (NR-mDec vs. NR-Dec: p < 0.01).

In regard to basal dendrites (Fig 3E, Suppl. Fig. 2E), we found a statistical general effect of behavior on the volume of stubby spines. Re-exposed mice had an increase of stubby spines’ volume compared to both non reexposed and behaviorally naïve animals (RE vs. NR: p < 0.05; RE vs. Na: p < 0.001). Furthermore, non re-exposed mice also had a larger volume than the behaviorally naïve ones (NR vs. Na: p < 0.01). We did not find a general effect of the drug in the measurement of this morphometric variable.

##### Thin spine volume

The volume of thin spines from apical dendrites (Fig 3C, Suppl. Fig. 2C) does not change either with behavior nor NF-κB inhibition. In basal dendrites (Fig 3F, Suppl. Fig. 2F), however, we found a significant general effect of behavior: animals that were re-exposed have smaller thin spines’ volume than non re-exposed animals (RE vs. NR: p < 0.001), and also compared to behaviorally naïve ones (RE vs. Na: p < 0.001).

#### (ii) Head diameter

##### Mushroom spine head diameter

When analyzing mushroom spines belonging to apical dendrites of animals (Fig 4A, Suppl. Fig. 3A) injected with the control solution mDec, we found that re-exposure increased their head diameter, compared to non re-exposed animals (RE-mDec vs. NR-mDec: p < 0.001). Inhibition of NF-κB prevented this increase (RE-mDec vs. RE-Dec: p < 0.01). Once again, we found an effect of NF-kB inhibition in non re-exposed animals. This time, the transcription factor’s inhibition increased the head diameter of mushroom spines (NR-mDec vs. NR-Dec: p < 0.001).

**Figure 4.**
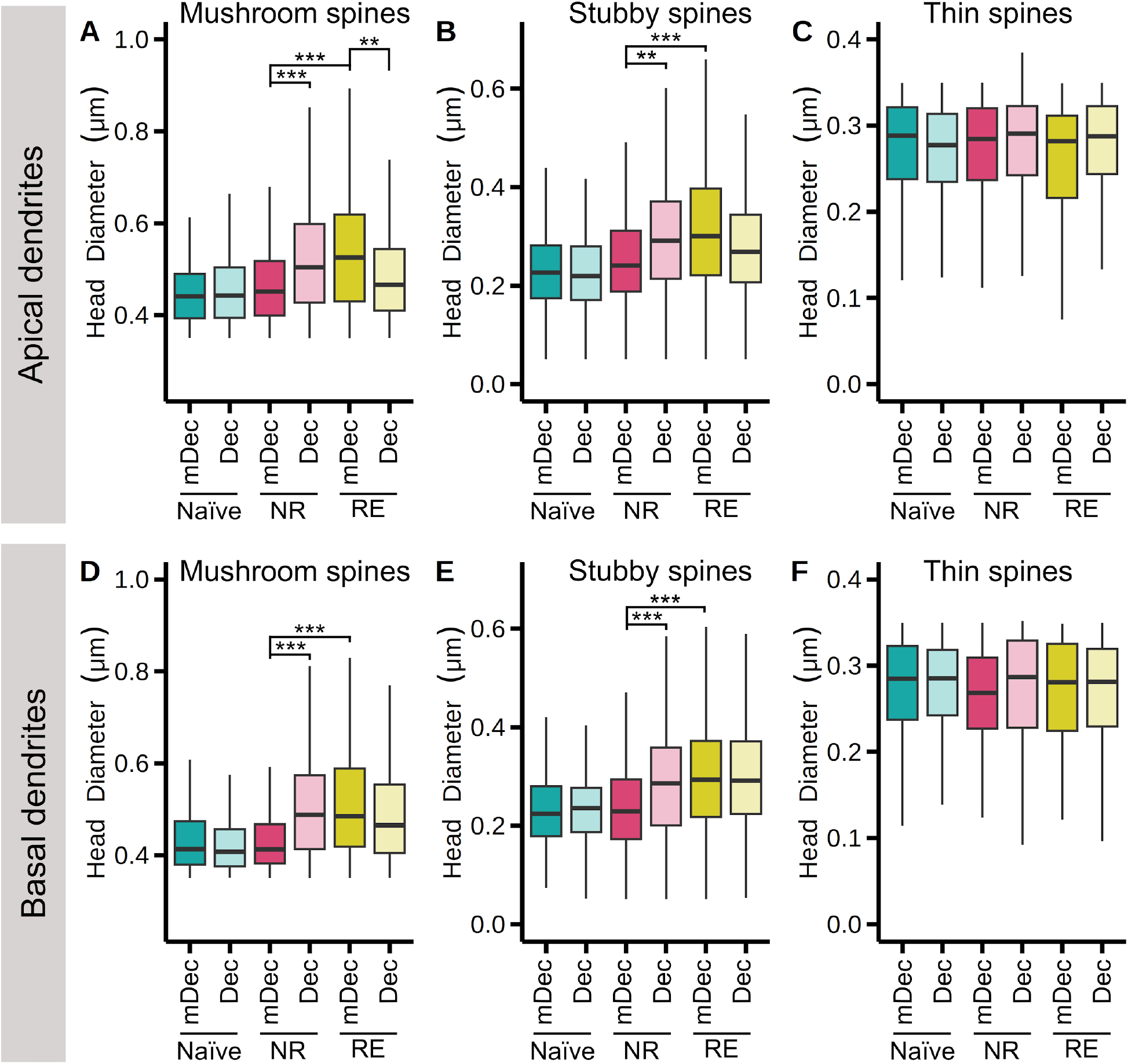
Long-term changes in head diameter of dendritic spines, analyzed per type of spine, associated with either the reconsolidation process or late-consolidation. Mushroom **(A)**, stubby **(B)** and thin **(C)** spine head diameter in the dorsal region of CA1, apical dendrites. **(D**,**E**,**F)** Same as in (A,B,C) in for basal dendrites. In all cases data are presented as box plots, depicting the first quartile, median, third quartile. The upper whisker extends to the largest value within 1.5 times the IQR, while the lower whisker extends to the smallest value within 1.5 times the IQR. n=2-5. ^**^P < 0.01, ^***^P < 0.001. Full statistical data can be found in Suppl. Table 2.

Similar results were obtained when analyzing head diameter of mushroom spines belonging to basal dendrites (Fig 4D, Suppl. Fig. 3D). However, in this case, NF-κB inhibition did not affect the head diameter in re-exposed animals (RE-mDec vs. RE-Dec: p = 0.446).

##### Stubby spine head diameter

Re-exposure increased the head diameter of stubby spines belonging to apical dendrites (Fig 4B, Suppl. Fig. 3B), compared to non re-exposed animals (RE-mDec vs. NR-mDec: p < 0.001). We again found an effect of NF-kB inhibition in non re-exposed animals. As happened with the head diameter of mushroom spines, NF-κB inhibition increased the head diameter of stubby spines (NR-mDec vs. NR-Dec: p < 0.001). Similar results were obtained when analyzing stubby spines from basal dendrites (Fig 4E, Suppl. Fig. 3E).

##### Thin spine head diameter

We did not find any statistical difference between groups in regard to the volume of mushroom spines, both in apical and basal dendrites (Fig 4C,F, Suppl. Fig. 3D,F).

#### iii) Neck diameter

Mushroom spine neck diameter. In apical dendrites (Fig 5A, Suppl. Fig. 4A) of animals injected with the control solution mDec, re-exposure increased the neck diameter of mushroom spines compared to non re-exposed animals (RE-mDec vs. NR-mDec: p < 0.001). Inhibition of NF-κB prevented this increase (RE-mDec vs. RE-Dec: p < 0.05). Once again, we found an effect of NF-kB inhibition in non re-exposed animals. In particular, NF-κB inhibition increased the neck diameter of mushroom spines (NR-mDec vs. NR-Dec: p < 0.001). Similar results were obtained when analyzing the neck diameter of mushroom spines belonging to basal dendrites (Fig 5C, Suppl. Fig. 4C).

**Figure 5.**
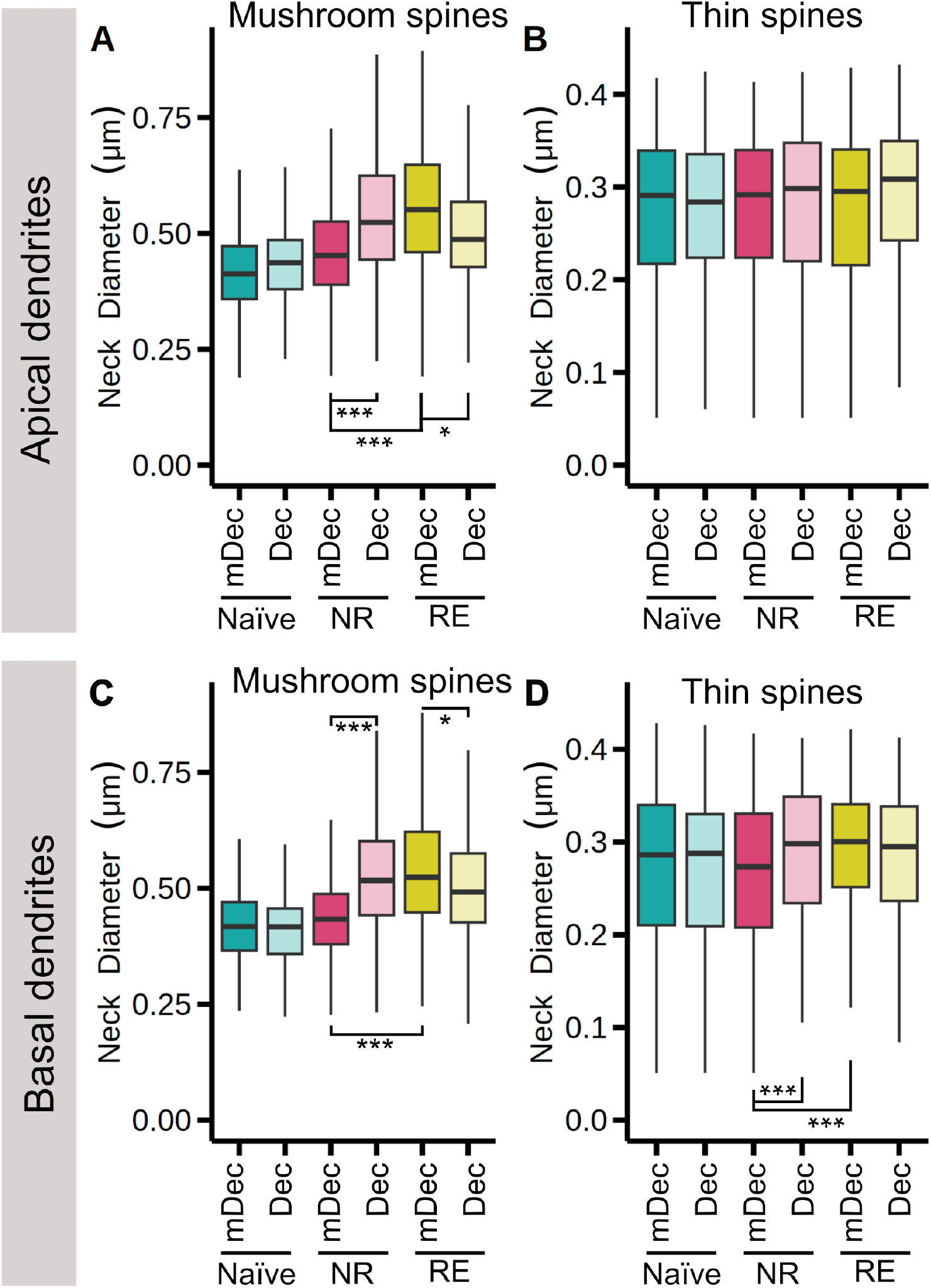
Long-term changes in neck diameter of dendritic spines, analyzed per type of spine, associated with either the reconsolidation process or late-consolidation. Mushroom **(A)**, stubby **(B)** and thin **(C)** spine neck diameter in the dorsal region of CA1, apical dendrites. **(D**,**E**,**F)** Same as in (A,B,C) in for basal dendrites. In all cases data are presented as box plots, depicting the first quartile, median, third quartile. The upper whisker extends to the largest value within 1.5 times the IQR, while the lower whisker extends to the smallest value within 1.5 times the IQR. n=2-5. ^*^P < 0.05, ^***^P < 0.001. Full statistical data can be found in Suppl. Table 2.

##### Thin spine neck diameter

In apical dendrites (Fig 5B, Suppl. Fig. 4B), we did not find any statistical difference between groups in regard to the neck diameter of thin spines. In turn, in basal dendrites (Fig 5D, Suppl. Fig. 4D) of animals injected with the control solution mDec, re-exposure increased neck diameter of mushroom spines compared to non re-exposed animals (RE-mDec vs. NR-mDec: p < 0.05). Here, as in the mushroom spines, we found an effect of NF-κB inhibition in non re-exposed animals: NF-κB inhibition increased the neck diameter of thin spines (NR-mDec vs. NR-Dec: p < 0.05).

It is relevant to mention that NF-κB inhibition did not affect the density of any type of spine, nor any morphometric parameter in behaviorally naïve animals, both in apical and basal dendrites (Table 1, Supplementary Table 1).

**Table 1.** Summary of the results obtained. Re-exposure effect column: *post hoc* comparison between RE-mDec and NR-mDec, or RE and NR in the case of simple effects. Training effect column: *post hoc* comparison between NR-mDec and Na-mDec, or NR and Na in the case of comparing main effects. RE column: *post hoc* comparison between RE-Dec and RE-mDec. NR column: *post hoc* comparison between NR-Dec and NR-mDec. Na column: *post hoc* comparison between Na-Dec and Na-mDec. Arrows up indicate an increase in the variable measured in the group in the left compared to the group in the right of each pair. Mushroom (orange) / Stubby (green) / Thin (red).

| Origin | Variable | Re-exposure effect | Training effect | NF-κ B's effect on | | |
| --- | --- | --- | --- | --- | --- | --- |
|  |  |  |  | RE | NR | Na |
| Apical | Density | — ↑ ↓ | — — — | ↓ — — | — — ↓ | — — — |
|  | Volume | — ↑ — | — — — | — ↓ — | — ↑ — | — — — |
|  | Head diameter | ↑ ↑ — | — — — | ↓ — — | ↑ ↑ — | — — — |
|  | Neck diameter | ↑ n/a — | — n/a — | ↓ n/a — | ↑ n/a — | — n/a — |
| Basal | Density | ↑ ↑ ↓ | — — ↓ | — — — | — — ↓ | — — — |
|  | Volume | — ↑ ↓ | — ↑ — | — — — | — — — | — — — |
|  | Head diameter | ↑ ↑ — | — — — | — — — | ↑ ↑ — | — — — |
|  | Neck diameter | ↑ n/a ↑ | — n/a — | ↓ n/a — | ↑ n/a ↑ | — n/a — |

## Discussion

In the present study we investigated dendritic spine changes associated with the reactivation of a contextual fear memory trace. In particular, we analyzed spine density and morphometric parameters of spines such as volume, head diameter and neck diameter. We specifically examined the spines located on the apical and basal dendrites of pyramidal neurons within the CA1 region of the dorsal hippocampus, a critical brain area associated with spatial and context-dependent associative memories^58–62^. Furthermore, we used NF-κB inhibition as a tool for negatively modulating memory reconsolidation, taking into account its effect at a behavioral level [Figure 1 and revised in ^42^] (Summary of results in Table 1).

We first found that re-exposure to the training context, a reminder that elicits the reactivation of the memory trace, does not alter long-term (*i.e*. 24h) total spine density in either of the sub-areas analyzed. However, when analyzing density per spine type, we did find changes. In particular, in apical dendrites of the CA1, we saw a marked decrease in the density of thin spines, together with an increase in density of stubby spines. These opposing effects could explain the lack of differences in total spine density. Similar results were obtained in basal dendrites, although in this case, re-exposure also increased mushroom spine density. In regard to the morphometric parameters analyzed, our results reveal that in apical dendrites re-exposure increases the volume of stubby spines, the head diameter of mushroom and stubby spines, and the neck diameter in mushroom spines. Similar results were observed in basal dendrites, where re-exposure also led to a decrease in the volume and an increase in the neck diameter of thin spines. Our results suggest that, at a population level, memory reactivation would generate a shift towards shapes that correlate with increased synaptic strength.

Importantly, inhibiting NF-κB prevented some, but not all, of the structural changes observed in animals that faced the reminder of the training event. In fact, NF-κB inhibition did not affect the decrease in volume of thin spines in basal dendrites, nor the increase in head diameter of stubby spines both in apical and basal dendrites, nor the increase in neck diameter of thin spines in basal dendrites. In turn, NF-κB inhibition indeed prevented the increase in neck diameter of mushroom spines, both from apical and basal dendrites. NF-κB inhibition prevented the increase in volume of stubby spines and the increase in head diameter of mushroom spines from apical, but not from basal dendrites. This differential effect of NF-κB can be explained by the fact that the NF-κB signaling pathway is indeed necessary for memory reconsolidation, but there are also other cellular pathways relevant for this process^63,64,27,65,66^. It makes sense that, at a behavioral level, blocking either of them results in a memory deficit. However, when focusing the analysis at other levels, such as structural plasticity, the differential contributions of each pathway might become evident. Our results suggest that NF-κB is not ubiquitously involved in all the structural changes associated with the reconsolidation process.

Many reconsolidation-eliciting protocols, such as the one used in this paper, include re-exposure to stimuli related to the learning event (*e.g*. the context). A frequently used control for this kind of studies consists of an experimental group where re-exposure is omitted (*i.e*. non re-exposed group), which usually serves to distinguish a reconsolidation-related process from a consolidation-derived one (such as persistence related mechanisms)^67,68^. On the one hand, our results demonstrate that, at a structural level, the non re-exposed control group differs from the behaviorally naïve control group, suggesting a long-term effect of consolidation itself. On the other hand, our work further reveals that, even when a drug does not have an effect on the measured behavioral output, it may have an impact on the brain. Indeed, NF-κB inhibition affected structural plasticity of dendritic spines in non re-exposed animals as well, compared to their respective control (non re-exposed animals, no NF-κB inhibition). The effects were observed in 10 out of 22 measurements (Table 1; Suppl. Table 1). To mention, NF-κB inhibition in non re-exposed animals diminished density of thin spines, both in apical and basal dendrites; increased the volume of stubby spines from apical dendrites; increased head diameter of mushroom and stubby spines, from apical and basal dendrites; and, finally, increased neck diameter of mushroom spines from apical and basal dendrites, and stubby spines from basal dendrites. Strikingly, inhibition of NF-κB in non-re-exposed animals had opposite effects to inhibition after re-exposure. More experiments are needed to understand this evidence in more detail.

Studies of neuronal structural plasticity in relation to memory reconsolidation are scarce^35–37^. In particular, one work aimed at studying structural changes in apical dendrites of CA1 of the dorsal hippocampus associated with memory reactivation at a long term^37^. Their approach differs from ours mainly in that their analysis and classification of spines is manual, and thus they measure density and not morphometric parameters. In particular, they found that reactivation induces a long-term increase in total spine density, which is also observed in non re-exposed animals, both compared to a naïve group. The authors suggest that the increase in spine density observed in reactivated animals is due to a reconsolidation-dependent process, as animals injected intra-amygdala with the drug propranolol, a blocker of reconsolidation, fail to show this increase. Interestingly, in non re-exposed animals, propranolol injection renders similar spine density than saline. Surprisingly, authors only discussed their findings centered on the increase due to reconsolidation but did not further discuss the plastic phenomena related to a long-term consolidation-dependent process observed in non re-exposed animals. As mentioned above, in our work, we also found that non re-exposed animals present a consolidation-dependent plasticity, not in spine density but indeed in other morphometric parameters. We believe that this plasticity which is unrelated to reactivation of the memory trace is remarkable, and that efforts should be made to better understand its role in memory persistence. Last, but not least, it is worth mentioning that neither their work nor ours evaluate the structural plasticity associated with reconsolidation in females, and this, at least in our case, is due to a carryover of past errors in which females were considered to have more variability than males^69^. In this respect, adult experimentally unmanipulated female rats present a cyclic fluctuation in spine density on apical dendrites from the CA1^70–72^, evidencing once more that results in males cannot be extrapolated to females. It would be absolutely relevant to study structural plasticity in females as well, not only in reconsolidation but in other phases of memory where females were understudied too.

Overall, our results support a model in which learning *per se* generates structural modifications that persist at least 48 h after the learning event, and for which some of them, but not all, needs NF-κB activation. In turn, the delivery of stimuli that induce labilization and reconsolidation also generates structural modifications, probably overlapping with those induced by the training session itself. Some of these reactivation-dependent modifications also rely on NF-κB activity. In the long term, total spine density does not change, but the density of each type of spine is indeed modified, favoring the presence of spines with morphologies that correlates with increased synaptic strength^3^. It has been shown that not all the neurons in a brain region that is relevant to the formation of a memory, indeed participate in the engram that encodes that memory ^(revised in 73,74)^. It would be very interesting to move forward with studies on neuronal morphology by focusing specifically on the neurons involved in the engram. Doing so represents a challenge, since the criteria used in studies aimed at identifying engram cells usually select these cells by the induction of the expression of certain genes, such as the immediate early gene cFOS^75,76^. These studies have demonstrated the necessity and sufficiency of these neurons, however, other neurons could also be involved in the memory trace and complementary studies would be useful to have a complete view of a very complex phenomenon. In turn, it would be very relevant to study whether reconsolidation induces spine clustering in these neurons that are part of the engram^2,4^.

## Methods

### Animals

Male mice, 8-12 weeks of age and weighing 25–30 g, were housed throughout the experimental procedures with water and food *ad libitum*, under a 12 h light/dark cycle (lights on at 8:00 A.M.) at a temperature of 21–23°C. For behavioral studies, we used C57BL/6 male mice from La Plata University animal facilities, La Plata, Argentina. For structural studies, we used male mice carrying the thy1-green fluorescent protein (GFP) transgene^77^ (Jackson Laboratory, Bar Harbor, ME, USA). Experiments were performed during the light cycle, and were designed and performed with the approval of the University of Buenos Aires Institutional Animal Care and Use Committee (CICUAL FCEN-UBA N°78/2017), and in accordance with regulations of the National Institutes of Health (NIH, USA) Guide for the Care and Use of Laboratory Animals (NIH publication 80-23/96). We made all efforts to minimize animal suffering and to reduce the number of animals used.

### Surgery and injections

Mice were bilaterally implanted under deep ketamine and xylazine anesthesia (100 mg/kg ketamine and 10 mg/kg xylazine, co-injected IP) with 23-gauge guide cannulae (coordinates: anterior-posterior -1.9 mm from bregma; medio-lateral ±1.2 mm; and dorso-ventral -1.2 mm from bregma, in accordance with the atlas of Franklin and Paxinos^78^ and personal adjustments^41^. While anesthetized, mice received one dose of analgesic (meloxicam, 5 mg/kg, SC). After surgeries, mice received analgesic (tramadol, 20 mg/ml, in drinking water for three days). Experiments were performed at least seven days after surgery to ensure animal recovery, and intra-hippocampal injections were administered without anesthesia after context reexposure session (or in the equivalent time point for non re-exposed and not trained animals). The injection device consisted of a 30-gauge cannula connected through a tubing to a 5 μL Hamilton syringe. The injection cannula was inserted into the guide cannula surpassing its length by 1 mm to reach the dorsal CA1 region of the hippocampus. The injections were manually administered during 30 s. The injection cannula was removed after 60 s to avoid reflux and to allow the diffusion of drugs.

### Drugs

As in de la Fuente et al., 2011^41^, NFκB Decoy (double-stranded DNA oligonucleotide 5’-GAG**GGGACTTTCC**CA-3’; consensus sequence in bold) and mDecoy (5’-GAG**G<u>C</u>GACTTTCC**CA-3’; base changed underlined)^46^ were dissolved in STE buffer solution. Decoy or mDecoy were used at a concentration of 0.47 g/L and delivered bilaterally 0.26 pmol per hemi-hippocampus at 0.5 μL per side, a dose that affects reconsolidation process^41^.

### Apparatus and behavioral procedures

The fear conditioning chamber was made of transparent acrylic and was located inside a wooden box. The floor of the chamber consisted of parallel stainless-steel grid bars through which we delivered foot shocks. Before training, animals were handled once a day for two days. In the training session, animals were placed individually in the conditioning chamber and allowed to acclimatize for two minutes. After this period, mice received three shock presentations (0.6 mA, 1 s, with an inter-trial interval of 1 min). Animals remained in the chamber for an additional minute and were returned to their home cages. A 5 min re-exposure to the training context performed 24 h after training was used to reactivate the memory trace (from here on “re-exposed”, RE)^41,49,51,79^. The testing session was performed 48 h after training (24 h post reactivation session). We used two behavioral controls: trained but non re-exposed animals (from here on “non re-exposed”, NR), and behaviorally naïve mice (littermates). Mice from each cage were assigned pseudorandomly to the experimental groups, trying to ensure that all groups were represented in all home cages.

All the sessions were video recorded for behavioral analysis. Memory was assessed in the re-exposure and testing sessions, and was expressed as the percentage of time that the mice spent freezing, a commonly used index of fear in mice, defined as the absence of all movements except for those related to breathing. Freezing was also measured in the training session, prior to the shocks, to establish baseline behavior. Freezing was scored manually by an observer blinded to the experimental condition, according to an instantaneous time sampling procedure in which each animal was observed every 5 s. We consider memory expression when the experimenters detect what has been specified as the operative measure of memory (in this case, freezing). Memory labilization comprises the circuital changes that makes the trace labile, and is elicited under the presentation of certain types of reminders. Reconsolidation, in turn, is the process that re-stabilizes that memory trace. In re-exposed animals injected with the control solution (mDecoy, mDec) we will not be able to differentiate between both processes, and thus the changes seen will be attributable to both. In pharmacological experiments in which the operative measure of memory is lower than in controls, we will assign the effect to a reconsolidation disruption (*e.g*. with Dec injection). The term reactivation will be used to refer to the circuital process that allows labilization/reconsolidation.

### Evaluation of density and morphological variables of dendritic spines

#### Brain fixation, sectioning and tissue preparation

Twenty-four hours after drug administration, animals were deeply anesthetized (ketamine and xylazine) and fixed by intracardiac perfusion of ∼10 mL of ice-cold 1% paraformaldehyde (PFA) in phosphate-buffer (PB), followed by ∼25 mL of ice-cold 4% PFA in PB. Brains were dissected, post-fixed overnight in the same fixing solution and maintained in PBS at 4 °C until used. Brains were sectioned in 130 μm slices, which were mounted using Vectashield (Vector Laboratories, Inc) over slides with SecureSeal™ imaging spacers (#654008, Grace Bio-Labs) and coverslips (474030-9020-000, Zeiss). Only data from animals with cannulae located in the intended sites were included in the analysis.

#### Acquisition of images

Apical (*stratum radiatum* and *lacunosum/moleculare*) and basal (*stratum oriens*) dendrites of the dorsal CA1 region of the hippocampus were imaged using an Olympus FV1000 confocal microscope and a 60 × objective (NA 1.42). Dendrites were chosen randomly within each *stratum*, excluding primary dendrites and prioritizing those most superficial to the sample. Total μm of apical dendrites measured per group: RE-Dec: 1304μm, RE-mDec: 842μm, NR-Dec: 1194μm, NR-mDec: 817μm, Na-Dec: 887μm, Na-mDec: 770μm. For basal dendrites: RE-Dec: 1063μm, RE-mDec: 755μm, NR-Dec: 774μm, NR-mDec: 695μm, Na-Dec: 821μm, Na-mDec: 816μm) Z-stacks spanning the entire thickness of each dendrite of interest were obtained at a resolution of 1024 × 1024 pixels. A linear 2× Kalman filter was used and, in order to obtain a 50 nm × 50 nm × 100 nm voxel size, images were acquired using a 4× digital zoom and 100 nm z-step.

### Density and morphological variables analyses using Neuronstudio

3D microscope Point-Spread Functions were generated from the confocal images using the Born & Wolf PSF optical model. Raw Images were deconvoluted for 10 iterations using the Richardson-Lucy algorithm (BatchDeconvolution^80^ plugin for ImageJ). The deconvoluted image was semi-automatically analyzed for individual dendritic spine identification using Neuronstudio^81,82^ (Computational Neurobiology and Imaging Center in ISMMS, Suppl. Table 2). Amount of spines in apical dendrites: RE-Dec: 2260, RE-mDec: 1815, NR-Dec: 2261, NR-mDec: 1869, Na-Dec: 2198, Na-mDec: 1642. In basal dendrites: RE-Dec: 1689, RE-mDec: 897, NR-Dec: 1195, NR-mDec: 1236, Na-Dec: 1627, Na-mDec: 1840.

#### Post-processing of dendritic and spine data

Spines were semi-automatically classified into three categories (stubby, thin and mushroom) according to the criteria stated on Table 1. The software output of the dendrite and spine analyses included the total number of spines, total length of the selected dendrite, spine volume, and head and neck (when applicable) diameter, among others. For total spine density analysis, the total amount of spines in each dendrite was divided by the total length of the dendrite. Spine density per type of spine was calculated similarly, but considering the amount of each type of spine per dendrite. Spine density per type of spine and morphometric parameters were calculated after excluding spines withneck’s diameter below the software detection limit (40% of total spines).

### Statistics

#### Behavioral data

Freezing scores were analyzed by General Linear Models using InfoStat (InfoStat 2016. Grupo InfoStat, FCA, Universidad Nacional de Córdoba, Argentina) and Akaike information criterion was considered in order to choose the best variance structure. Fisher’s LSD was used for *post hoc* comparisons and the Š*idák* correction method was used to adjust for multiple comparisons.

#### Spine density and morphological variables

Data analyses were performed using R (version 4.2.2,^83^). To account for data dependency (as multiple observations were drawn from each mouse), a generalized linear mixed model (GLMM) was performed using the glmmTMB package^84^ (version 1.1.6). We fitted a GLMM for each morphological variable of interest with Drug (mDec or Dec) and Behavior (RE, NR, Na) as fixed effects and Mouse ID for random effects. The GLMMs corresponding to the variables head diameter, neck diameter and volume were fitted with Gamma family distribution and a log link function, while the GLMM for spine density was fitted with Gaussian family distribution and a log link function. The goodness-of-fit for each model was analyzed using the DHARMa package^85^ (version 0.4.6). Visual inspection of residual plots did not reveal any concerning deviations from normality or homoscedasticity. Post-hoc comparisons were performed using the emmeans package^86^ (version 1.8.5) and all reported p-values were adjusted for multiple comparisons with the Šidák correction method.

## Acknowledgements

The research was supported by grants to VdlF from the Agencia Nacional de Promoción Científica y Tecnológica of Argentina (PICT 2017, 2019) and CAEN grant from the International Society for Neurochemistry, and grants to AR from the Agencia Nacional de Promoción Científica y Tecnológica of Argentina (PICT 2015, 2018). We are indebted to IBRO for funding VdlF visit to Pozzo Miller Lab at University of Alabama at Birmingham, and Fulbright-BEC.AR for funding VdlF to visit Dani Dumitriu Lab at Mount Sinai Hospital, NY. This work was supported by grants from the Argentine Agency for the Promotion of Science and Technology

## Additional Information (including a Competing Interests Statement)

Authors declare no competing interests

