## Supporting material for "Long-term structural plasticity of hippocampal dendritic spines following contextual fear memory reactivation"

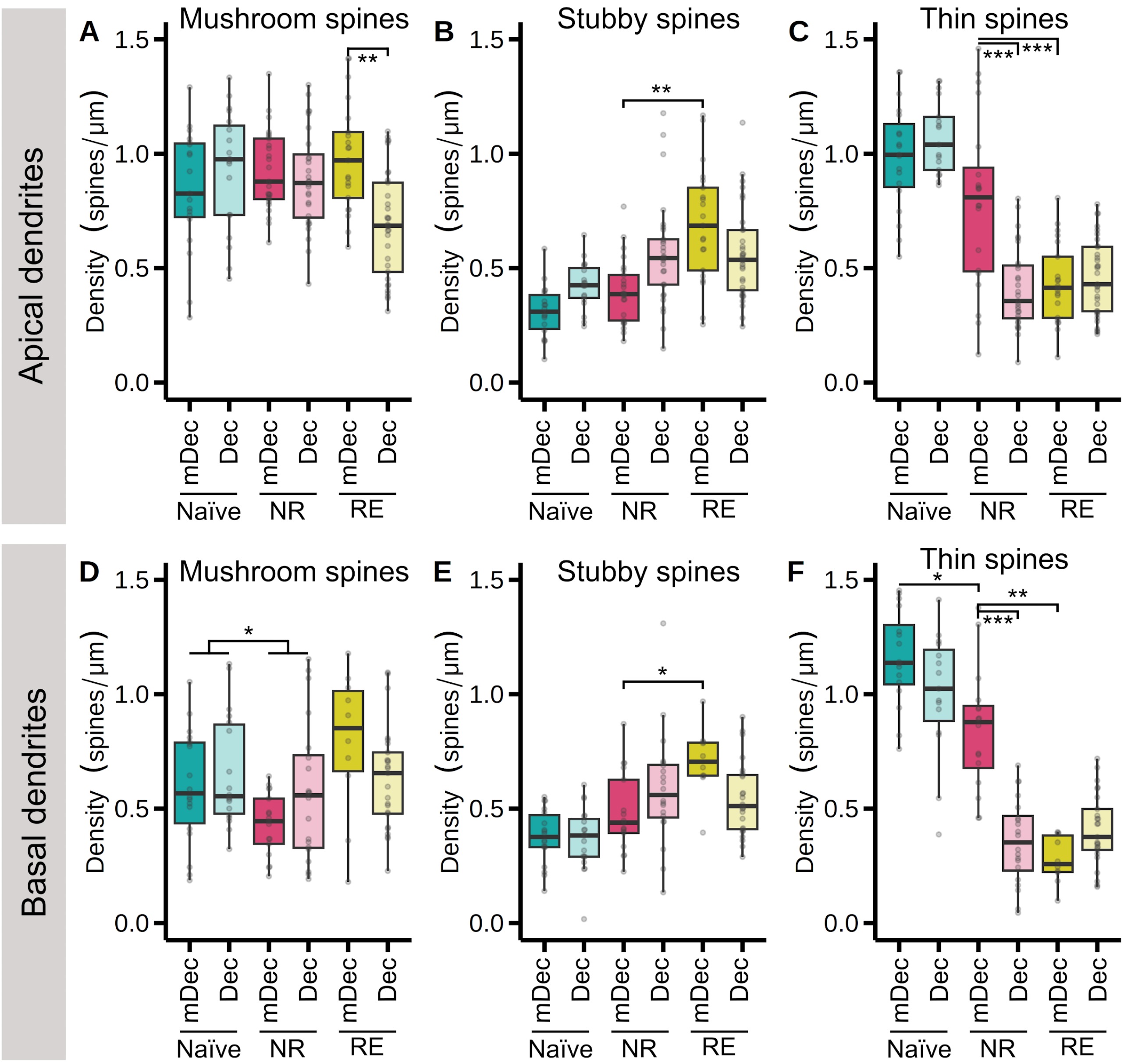


Supplementary Figure 1. **Long-term changes in spine density, analyzed per type of spine, associated with either the reconsolidation process or late-consolidation.** Mushroom **(A)**, stubby **(B)** and thin **(C)** spine density of dorsal CA1 apical dendrites. **(D,E,F)** Same as in (A,B,C) basal dendrites. In all cases data are presented as box plots, depicting the first quartile, median, third quartile. The upper whisker extends to the largest value within 1.5 times the IQR, while the lower whisker extends to the smallest value within 1.5 times the IQR. Dots represent values of individual spines. n=2-5. *P < 0.05, **P < 0.01, ***P < 0.001. Full statistical data can be found in Suppl. Table 2.

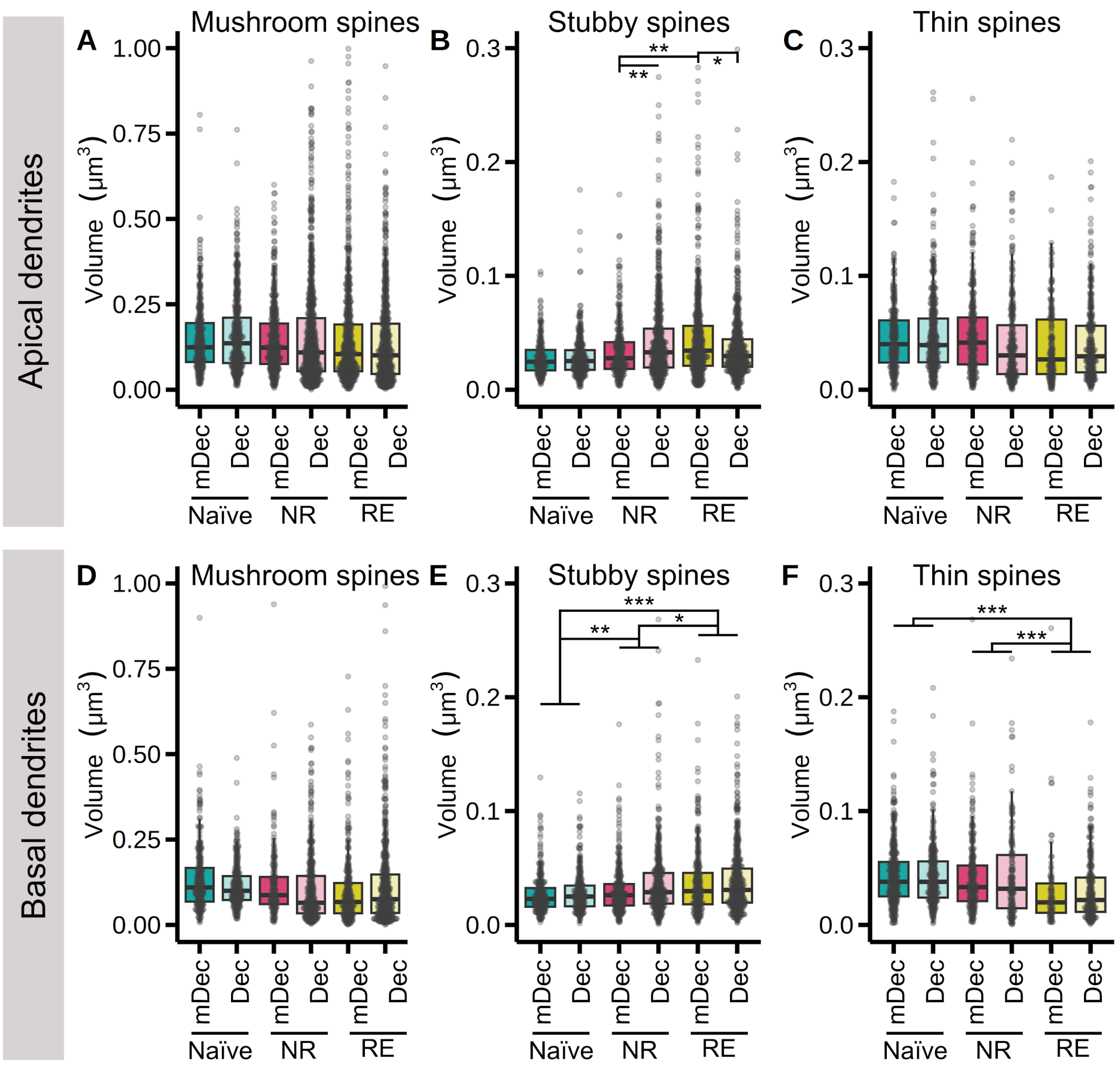


Supplementary Figure 2. **Long-term changes in volume of dendritic spines, analyzed per type of spine, associated with either the reconsolidation process or late-consolidation.** Mushroom **(A)**, stubby **(B)** and thin **(C)** spine volume in the dorsal region of CA1, apical dendrites. **(D,E,F)** Same as in (A,B,C) in for basal dendrites. In all cases data are presented as box plots, depicting the first quartile, median, third quartile. The upper whisker extends to the largest value within 1.5 times the IQR, while the lower whisker extends to the smallest value within 1.5 times the IQR. Dots represent values of individual spines. n=2-5. *$P < 0.05$, **$P < 0.01$, ***$P < 0.001$. Full statistical data can be found in Suppl. Table 2.

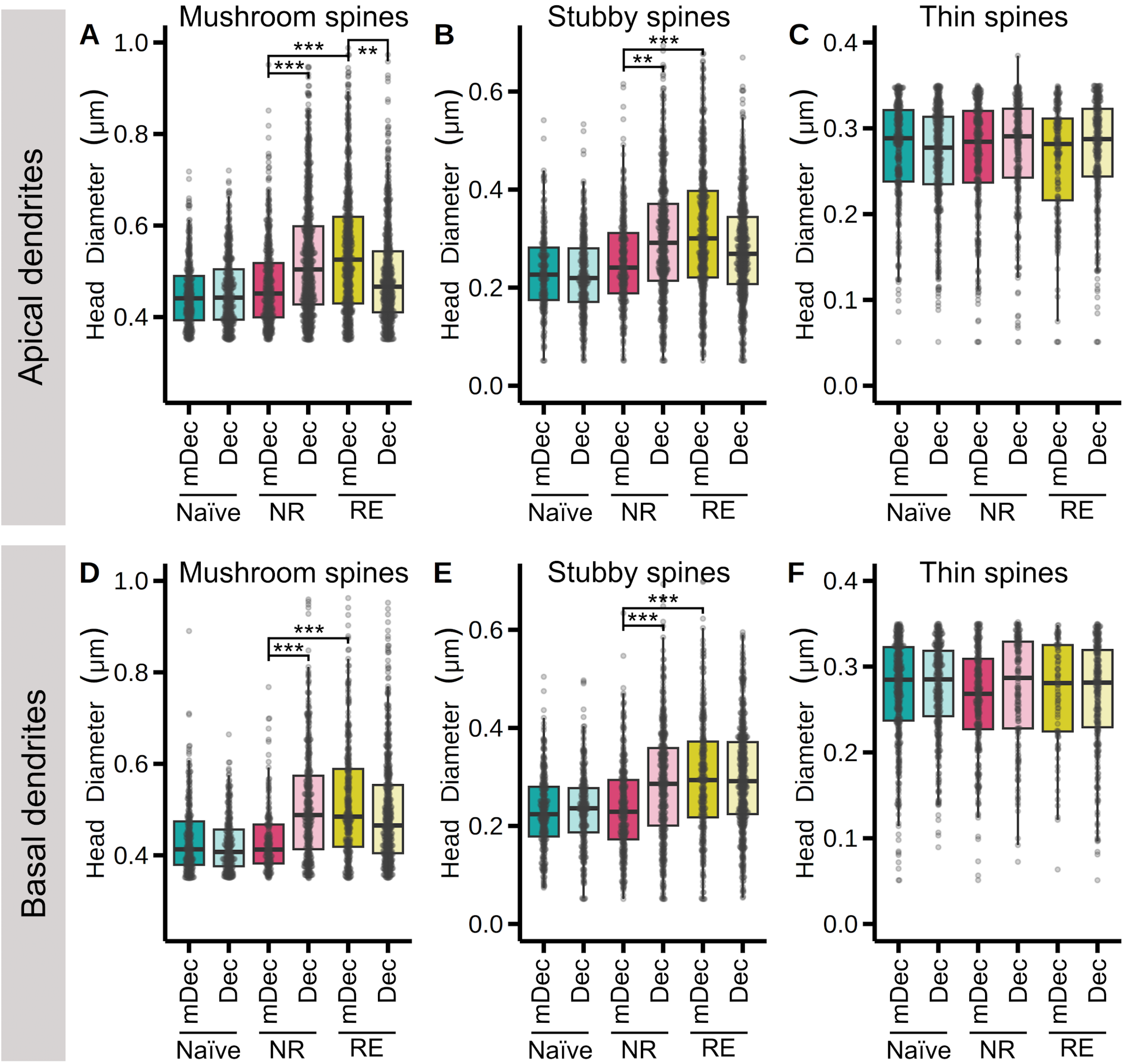


Supplementary Figure 3. **Long-term changes in head diameter of dendritic spines, analyzed per type of spine, associated with either the reconsolidation process or late-consolidation.** Mushroom **(A)**, stubby **(B)** and thin **(C)** spine head diameter in the dorsal region of CA1, apical dendrites. **(D,E,F)** Same as in (A,B,C) in for basal dendrites. In all cases data are presented as box plots, depicting the first quartile, median, third quartile. The upper whisker extends to the largest value within 1.5 times the IQR, while the lower whisker extends to the smallest value within 1.5 times the IQR. Dots represent values of individual spines. n=2-5. $^{**}P < 0.01$, $^{***}P < 0.001$. Full statistical data can be found in Suppl. Table 2.

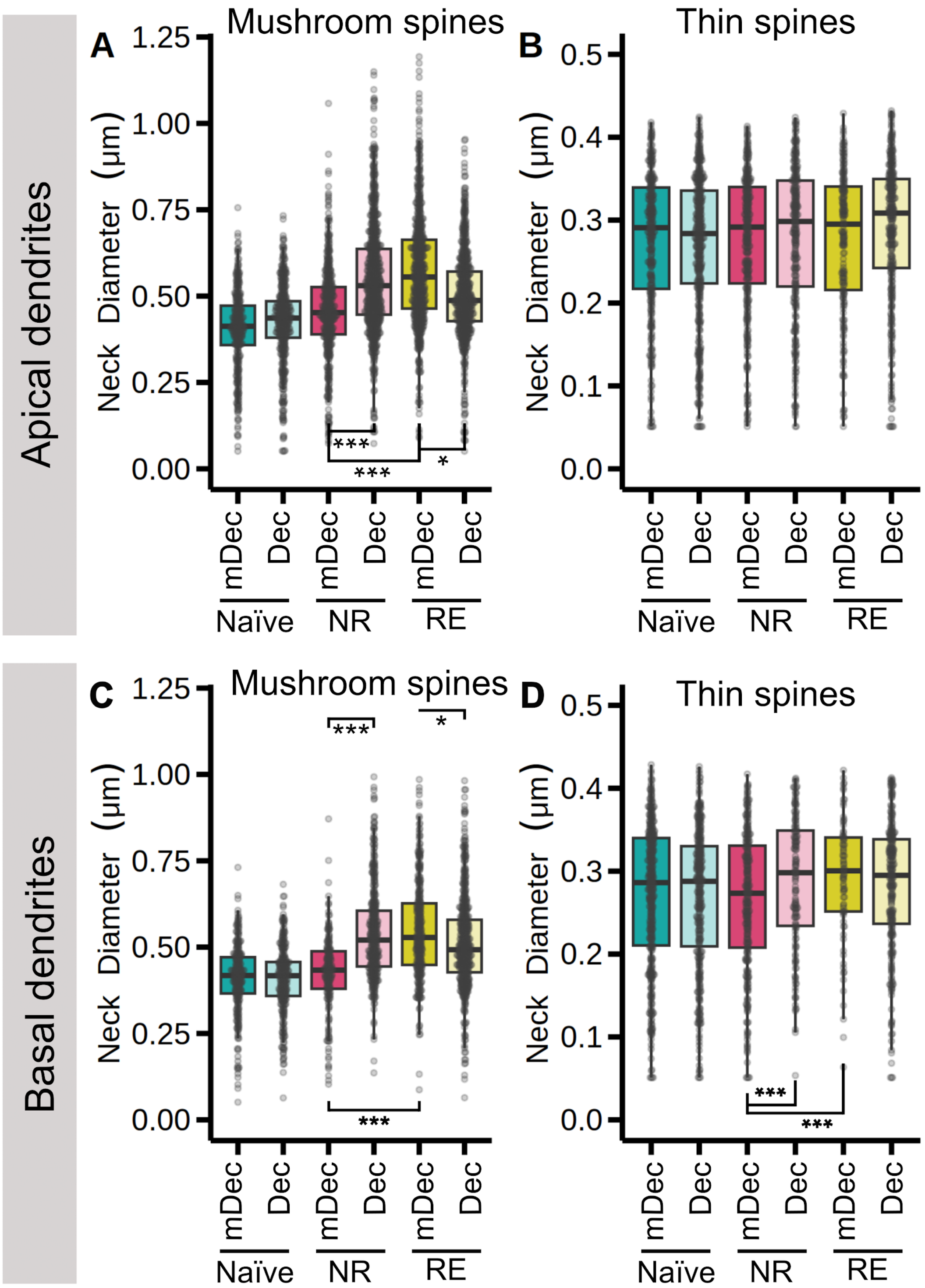


Supplementary Figure 4. **Long-term changes in neck diameter of dendritic spines, analyzed per type of spine, associated with either the reconsolidation process or late-consolidation.** Mushroom **(A)**, stubby **(B)** and thin **(C)** spine neck diameter in the dorsal region of CA1, apical dendrites. **(D,E,F)** Same as in (A,B,C) in for basal dendrites. In all cases data are presented as box plots, depicting the first quartile, median, third quartile. The upper whisker extends to the largest value within 1.5 times the IQR, while the lower whisker extends to the smallest value within 1.5 times the IQR. Dots represent values of individual spines. n=2-5. *P < 0.05, ***P < 0.001. Full statistical data can be found in Suppl. Table 2.

**<u>Supplementary Table 1</u>**. Results of the statistical analyses (the effect of NF-κB inhibition in non re-exposed animals is depicted in pink)

| **Supplementary Table 1** | | | | | | |
|---|---|---|---|---|---|---|
| **Variable** | **Fig. ref.** | | **Group Effect** | **Session Effect** | **Interaction** | **Contrast** |
| Behavior | 1C | | **df = 1**<br>**F = 7.402**<br>**p = 0.015** | **df = 2**<br>**F = 168.4**<br>**p = 1.0e-04** | **df = 1**<br>**F = 9.645**<br>**p = 5.0e-04** | RE: p = 0.7618<br>**TS: p = 3.0e-04** |
| Behavior | 1F | | df = 1<br>F = 1.362<br>p = 0.29 | **df = 1**<br>**F = 81.56**<br>**p = 1.0e-04** | df = 1<br>F = 0.8676<br>p = 0.39 | - |
| **Variable** | **Location** | **Type** | **Drug Effect (df = 1)** | **Behavior Effect (df = 2)** | **Interaction (df = 2)** | **Contrast** |
| Spine Density | Apical | - | F = 3.563<br>p = 0.059 | **F = 9.193**<br>**p = 0.010** | **F = 9.162**<br>**p = 0.010** | RE-mDec - RE-Dec: p = 0.092<br>NR-mDec - NR-Dec: p = 0.14<br>RE-mDec - NR-mDec: p = 0.93<br>Na-mDec - Na-Dec: p = 0.50<br>Na-mDec - NR-mDec: p = 0.94 |
| | Basal | - | **F = 3.248**<br>**p = 0.072** | **F = 11.60**<br>**p = 0.0030** | F = 0.4991<br>p = 0.78 | RE - NR: p = 0.99<br>**RE - Na: p = 0.029**<br>**NR - Na: p = 0.0074** |
| Spine Density | Apical | Mushroom | F = 2.949<br>p = 0.086 | F = 2.975<br>p = 0.22 | **11.45**<br>**p = 0.0033** | **RE-mDec - RE-Dec: p = 0.0017**<br>NR-mDec - NR-Dec: p = 1.0<br>RE-mDec - NR-mDec: p = 0.96<br>Na-mDec - Na-Dec: p = 0.88<br>Na-mDec - NR-mDec: p = 0.89 |
| | | Stubby | F = 1.261<br>p = 0.26 | **F = 16.91**<br>**p = 2.1e-04** | **F = 9.231**<br>**p = 0.0099** | RE-mDec - RE-Dec: p = 0.48<br>NR-mDec - NR-Dec: p = 0.095<br>**RE-mDec - NR-mDec: p = 0.0027**<br>Na-mDec - Na-Dec: p = 0.45<br>Na-mDec - NR-mDec: p = 0.67 |
| | | Thin | **F = 5.050**<br>**p = 0.025** | **F = 40.03**<br>**p = 2.0e-09** | **F = 17.76**<br>**p = 1.4e-04** | RE-mDec - RE-Dec: p = 1.0<br>**NR-mDec - NR-Dec: p = 3.0e-05**<br>**RE-mDec - NR-mDec: p = 4.5e-04**<br>Na-mDec - Na-Dec: p = 0.95<br>Na-mDec - NR-mDec: p = 1.0 |
| | Basal | Mushroom | F = 0.07302<br>p = 0.79 | **F = 6.194**<br>**p = 0.045** | F = 3.455<br>p = 0.18 | **RE - NR: p = 0.016**<br>RE - Na: p = 0.71<br>NR - Na: p = 0.14 |
| | | Stubby | F = 0.5302<br>p = 0.47 | **F = 25.61**<br>**p = 2.8e-06** | **F = 6.904**<br>**p = 0.032** | RE-mDec - RE-Dec: p = 0.10<br>NR-mDec - NR-Dec: p = 0.56<br>**RE-mDec - NR-mDec: p = 0.018**<br>Na-mDec - Na-Dec: p = 1.0<br>Na-mDec - NR-mDec: p = 0.37 |
| | | Thin | **F = 8.526**<br>**p = 0.0035** | **F = 70.10**<br>**p = 6.0e-16** | **F = 16.61**<br>**p = 2.5e-04** | RE-mDec - RE-Dec: p = 0.76<br>**NR-mDec - NR-Dec: p = 3.3e-05**<br>**RE-mDec - NR-mDec: p = 0.0018**<br>Na-mDec - Na-Dec: p = 0.81<br>**Na-mDec - NR-mDec: p = 0.036** |
| Spine Volume | Apical | Mushroom | F = 0.001621<br>p = 0.97 | F = 1.032<br>p = 0.60 | F = 2.724<br>p = 0.26 | - |
| | | Stubby | F = 0.6688<br>p = 0.41 | **F = 40.07**<br>**p = 2.0e-09** | **F = 19.72**<br>**p = 5.2e-05** | **RE-mDec - RE-Dec: p = 0.048**<br>**NR-mDec - NR-Dec: p = 0.0012**<br>**RE-mDec - NR-mDec: p = 0.0018**<br>Na-mDec - Na-Dec: p = 1.0<br>Na-mDec - NR-mDec: p = 0.17 |
| | | Thin | F = 0.3691<br>p = 0.54 | F = 3.056<br>p = 0.22 | F = 4.633<br>p = 0.099 | - |

| | | | | | | |
|---|---|---|---|---|---|---|
| | Basal | Mushroom | F = 0.03108<br>p = 0.86 | F = 2.734<br>p = 0.25 | F = 5.632<br>p = 0.060 | - |
| | | Stubby | F = 3.162<br>p = 0.075 | **F = 31.73**<br>**p = 1.3e-07** | F = 3.076<br>p = 0.21 | **RE - NR: p = 0.014**<br>**RE - Na: p = 2.0e-08**<br>**NR - Na: p = 0.0044** |
| | | Thin | F = 0.4020<br>p = 0.53 | **F = 27.99**<br>**p = 8.3e-07** | F = 0.2010<br>p = 0.90 | **RE - NR: p = 5.2e-08**<br>**RE - Na: p = 2.4e-12**<br>NR - Na: p = 0.55 |
| Head Diameter | Apical | Mushroom | F = 0.8462<br>p = 0.36 | **F = 33.71**<br>**p = 4.8e-08** | **F = 32.01**<br>**p = 1.1e-07** | **RE-mDec - RE-Dec: p = 0.0030**<br>**NR-mDec - NR-Dec: p = 2.4e-05**<br>**RE-mDec - NR-mDec: p = 1.6e-05**<br>Na-mDec - Na-Dec: p = 1.0<br>Na-mDec - NR-mDec: p = 0.60 |
| | | Stubby | F = 0.2690<br>p = 0.60 | **F = 46.34**<br>**p = 8.7e-11** | **F = 18.08**<br>**p = 1.2e-04** | RE-mDec-RE-Dec: p = 0.090<br>**NR-mDec-NR-Dec: p = 0.0021**<br>**RE-mDec-NR-mDec: p = 5.7e-04**<br>Na-mDec-Na-Dec: p = 1.0<br>Na-mDec-NR-mDec: p = 0.38 |
| | | Thin | F = 0.01783<br>p = 0.89 | F = 1.207<br>p = 0.55 | F = 4.299<br>p = 0.12 | - |
| | Basal | Mushroom | F = 2.886<br>p = 0.089 | **F = 45.56**<br>**p = 1.3e-10** | **F = 27.66**<br>**p = 9.9e-07** | RE-mDec - RE-Dec: p = 0.45<br>**NR-mDec - NR-Dec: p = 8.40e-07**<br>**RE-mDec - NR-mDec: p = 1.3e-05**<br>Na-mDec - Na-Dec: 0.97<br>Na-mDec - NR-mDec: p = 1.0 |
| | | Stubby | **F = 9.158**<br>**p = 0.0025** | **F = 39.08**<br>**p = 3.3e-09** | **F = 8.997**<br>**p = 0.011** | RE-mDec - RE-Dec: p = 1.0<br>**NR-mDec - NR-Dec: p = 1.5e-04**<br>**RE-mDec - NR-mDec: p = 2.0e-04**<br>Na-mDec - Na-Dec: p = 0.92<br>Na-mDec - NR-mDec: p = 0.98 |
| | | Thin | F = 0.8279<br>p = 0.36 | F = 5.300<br>p = 0.071 | F = 2.373<br>p = 0.30 | - |
| Neck Diameter | Apical | Mushroom | F = 2.668<br>p = 0.10 | **F = 56.39**<br>**p = 5.7e-13** | **F = 29.02**<br>**p = 5.0e-07** | **RE-mDec - RE-Dec: p = 0.021**<br>**NR-mDec - NR-Dec: p = 9.9e-06**<br>**RE-mDec - NR-mDec: p = 3.0e-06**<br>Na-mDec - Na-Dec: p = 0.91<br>Na-mDec - NR-mDec: p = 0.092 |
| | | Thin | F = 0.5190<br>p = 0.47 | F = 3.154<br>p = 0.21 | F = 1.184<br>p = 0.55 | - |
| | Basal | Mushroom | **F = 7.822**<br>**p = 0.0052** | **F = 181.4**<br>**p = 4.1e-40** | **F = 71.80**<br>**p = 2.6e-16** | **RE-mDec - RE-Dec: p = 0.020**<br>**NR-mDec - NR-Dec: p = 3.2e-23**<br>**RE-mDec - NR-mDec: p = 5.6e-16**<br>Na-mDec - Na-Dec: p = 1.0<br>Na-mDec - NR-mDec:p = 0.73 |
| | | Thin | F = 0.4843<br>p = 0.49 | F = 5.636<br>p = 0.060 | **F = 7.815**<br>**p = 0.020** | RE-mDec - RE-Dec: p = 0.91<br>**NR-mDec - NR-Dec: p = 0.034**<br>**RE-mDec - NR-mDec: p = 0.044**<br>**Na-mDec - Na-Dec: p = 0.0030**<br>Na-mDec - NR-mDec: p = 0.78 |

**<u>Supplementary Table 2</u>**. Value of the identification parameters and classification criteria used for semi-automatic detection and classification of dendrites and spines using Neuronstudio software.

| **Identification parameters** | | | **Value** |
|---|---|---|---|
| Residual Smear | | | medium (1.5x) |
| Neurite min. length | | | 5 µm |
| Spines | | Min. height | 0.2 µm |
| | | Max. height | 3 µm |
| | | Max. width | 3 µm |
| | | Min. non-stubby size | 5 vox |
| | | Min. stubby size | 5 vox |
| | | Correction | 25% |
| Spine classifier | | Neck ratio | 0.8 |
| | | Thin ratio | 2 |
| | | Mushroom size | 0.35 |
| **Classification criteria** | **Spine type** | | |
| | thin | mushroom | stubby |
| head diameter/neck diameter > 0.8 | ✓ | ✓ | × |
| spine length/head diameter > 2 | ✓ | × | × |
| head diameter > 0.35 µm | × | ✓ | × |
